# γ-H2AX is present at mouse meiotic kinetochores

**DOI:** 10.1101/2020.03.10.986273

**Authors:** Andrea Guajardo, Alberto Viera, María Teresa Parra, Manuel M. Valdivia, Julio S. Rufas, José A. Suja

## Abstract

The histone variant H2AX phosphorylated on serine 139, named γ-H2AX, is a canonical DNA double-strand breaks marker. During mammalian meiotic prophase I, γ-H2AX participates in meiotic recombination, meiotic sex chromosome inactivation and meiotic silencing of unsynapsed chromatin. In this study, we have analyzed the distribution of γ-H2AX during male mouse meiosis by immunofluorescence on spread and squashed spermatocytes. We have found that γ-H2AX locates at the inner kinetochore plate of meiotic kinetochores in both meiotic divisions. Therefore our results, for the first time, uncover a novel role for γ-H2AX at mammalian meiotic kinetochores.

## Introduction

Post-translational histone modifications and histone variants are involved in somatic cells in regulating chromatin dynamics that affect, among other processes, to DNA replication and repair, transcription and chromosome packaging and segregation (Tessarz and Kouzarides, 2014; Talbert and Henikoff, 2017). Likewise, during meiosis, a specialized cell division process that produces haploid gametes by performing two consecutive rounds of chromosome segregation after a single round of DNA replication (Bolcun-Filas and Handel, 2018), a “meiotic histone code” regulates chromatin remodeling, DNA double-strand breaks (DSBs) formation and repair, synapsis, recombination, meiotic sex chromosome inactivation, transcription, meiosis resumption and chromosome condensation and segregation (Wang et al., 2017).

Histone H2AX is a histone H2A variant that is characterized by a unique motif in its C-terminal domain (Millar, 2013). Phosphorylation of histone H2AX on serine 139 by phosphatidylinositol 3-kinase protein kinase-like (PIKK) members ATM, ATR and DNA-PK is called γ-H2AX (Durocher and Jackson, 2001). γ-H2AX plays an important canonical role in the DNA damage response (DDR) located at the chromatin surrounding DSBs, so dysfunctions of γ-H2AX lead to genomic instability (Georgoulis et al., 2017). In addition, it has been suggested that during mitosis the phosphorylation of H2AX by ATM may contribute to the fidelity of the mitotic process in spite of the absence of DNA damage to maintain genome integrity (McManus and Hendzel, 2005; Ichijima et al., 2005). In this regard, it has been proposed that during mitosis the kinase ATM phosphorylates histone H2AX on serine 139 at kinetochores, and that γ-H2AX recruits the DDR protein MDC1 that in turn regulates the loading of the spindle-assembly checkpoint (SAC) proteins MAD2 and CDC20 to the kinetochores (Eliezer et al., 2014). The SAC regulates metaphase to anaphase transition preventing entry into anaphase until all sister kinetochores are attached to the microtubules from opposite spindle poles ensuring the fidelity of chromosome segregation (Lara-Gonzalez et al., 2012; Dou et al., 2019). Several SAC components such as MAD1, MAD2, MAD3/BUBR1, BUB1, and BUB3 are localized to unattached kinetochores and participate in a signaling network that leads the inhibition of the E3 ubiquitin ligase anaphase promoting complex/cyclosome (APC/C) through recruiting CDC20, an activation factor of the APC/C (Kapanidou et al., 2017).

The DDR kinase ATM also phosphorylates BUB1 and MAD1 in the absence of DNA damage, and these phosphorylations are essential for the activation of the SAC during mitosis (Yang et al., 2011; Yang et al., 2014). In addition, the DDR protein 53BP1 also localizes to kinetochores and is hyperphosphorylated when the SAC is activated during mitosis, which has allowed to propose that 53BP1 has a role in kinetochore signaling in addition to its role in DDR (Jullien et al., 2002). Interestingly, other canonical DDR proteins such as the kinase ATR, CHK1, and RPA (Kabeche et al., 2018), as well as CHK2 (Petsalaki and Zachos, 2014) and BRCA2 (Park et al., 2017), also localize to mitotic kinetochores. Thus, several evidences point to an important link between the DDR and the SAC, the two major checkpoints that maintain the genome integrity (Lawrence and Engebrecht, 2015; Petsalaki and Zachos, 2020).

The dynamics of γ-H2AX has been extensively analyzed during mouse prophase I stages. During the leptotene stage of prophase I meiotic recombination initiates in mammals by the formation of programmed DSBs by the endonuclease SPO11 (Yamada et al., 2017), which leads to the appearance of γ-H2AX in the surrounding chromatin (Mahadevaiah et al., 2001). γ-H2AX is involved in the recruitment and activation of DDR proteins like MDC1, BRCA1, 53BP1 and RAD51 at sites of DSBs (Wang et al., 2017). In addition, γ-H2AX has also been linked with meiotic sex chromosome inactivation (MSCI), a process by which transcription of the X and Y chromosomes is repressed at the sex body (Fernandez-Capetillo et al., 2003; Turner et al., 2004; Turinetto and Giachino, 2015). Moreover, γ-H2AX is also involved in the meiotic silencing of unsynapsed chromosomes (MSUC) (Baarends et al., 2005; Turner et al., 2005; Manterola et al., 2009). It has been reported that deletion of histone H2AX in mice results in male infertility and pachytene arrest which are associated with defects in the pairing of the sex chromosomes (Celeste et al., 2002). However, it has also been highlighted that the presence of γ-H2AX labeling is not restricted to prophase I spermatocytes, since it labels chromatin in some spermatogonia and spermatids in mouse (Hamer et al., 2003). In this sense, it has been described that the spermatid kinase TSSK6 is essential for fertility of male mice mediating γ-H2AX formation that is required for a correct histone-to-protamine transition during spermiogenesis (Jha et al., 2017). Altogether, these data demonstrate that γ-H2AX is involved in different biological processes during mouse spermatogenesis.

Although significant progress in γ-H2AX function during mammalian meiosis has been made, all studies performed so far mainly focused on prophase I spermatocytes and spermatids. Thus, γ-H2AX behavior during mammalian meiotic divisions has never been examined. Here, using several antibodies against γ-H2AX we found that this posttranslational modification appears at kinetochores during both mouse meiotic divisions suggesting that, in addition to its role in DDR, it may have a role at kinetochores during meiosis.

## Results

### Post-diplotene distribution of γ-H2AX

We first analyzed by immunofluorescence the distribution of γ-H2AX on spread mouse spermatocytes using three different specific antibodies for this histone variant commonly used in mouse spermatocytes and for SYCP3, a structural component of the axial/lateral elements of the synaptonemal complex, to identify the different meiotic stages. The three antibodies against γ-H2AX used in this study offered the same staining patterns. As previously reported (Mahadevaiah et al., 2001), γ-H2AX labeling covered most of the chromatin during leptotene when the axial elements (AEs) begin to assemble along the chromosomes (Fig. S1A-D). In early zygotene spermatocytes, when AEs start to synapse and are then called lateral elements (LEs), which were visualized as thicker filaments of SYCP3, the pattern of γ-H2AX was similar to that found in the leptotene stage. As the zygotene stage progressed, the γ-H2AX labeling started to decrease and mostly surrounded chromatin from unsynapsed chromosome regions (Fig. S1E and F). In late zygotene spermatocytes, when only the sex chromosomes remained unsynapsed, most γ-H2AX labeling accumulated over the chromatin of these chromosomes (Fig. S1G and H). During pachytene and up to a late diplotene stage, most of the γ-H2AX signal was found at the sex body (Fig. S1I-L).

During diakinesis the γ-H2AX labeling was also restricted to the chromatin of sex chromosomes, showing a more discontinuous and diffuse pattern than that previously visualized during the pachytene and diplotene stages (Fig. 1A-C). The γ-H2AX labeling on the sex chromatin was still evident in all prometaphase I spermatocytes (Fig. 1D-F), and in 52% (n=42) of metaphase I ones (Fig. 1G-I; left top inset in H). Surprisingly, in prometaphase I and metaphase I spermatocytes γ-H2AX also appeared as pairs of closely associated dots (Fig. 1D-I) located at the chromatin regions more brightly stained with DAPI which correspond to centromeric regions (Fig. 1F and I). During metaphase I, SYCP3 was observed accumulated at the inner centromere domain (Parra et al., 2004; Gómez et al., 2007) appearing as T-shaped signals in side-viewed centromeres (Fig. 1H; right top inset) or as two side-by-side associated rings when centromeres were top-viewed (Fig. 1H; right bottom inset). During this stage, small SYCP3 patches appeared at the so-called interchromatid domain along the surface of contact between sister chromatid arms (Fig. 1I; Parra et al., 2004). In side-viewed centromeres, γ-H2AX was present as a pair of closely associated dots above the T-shaped SYCP3 signal (Fig. 1H; right top inset) or encircled by the SYCP3 rings when centromeres were top-viewed (Fig. 1H; right bottom inset). This distribution pattern of γ-H2AX is reminiscent of other kinetochore proteins previously described at metaphase I centromeres (Parra et al., 2004; Gomez et al., 2007).

**Fig. 1.**
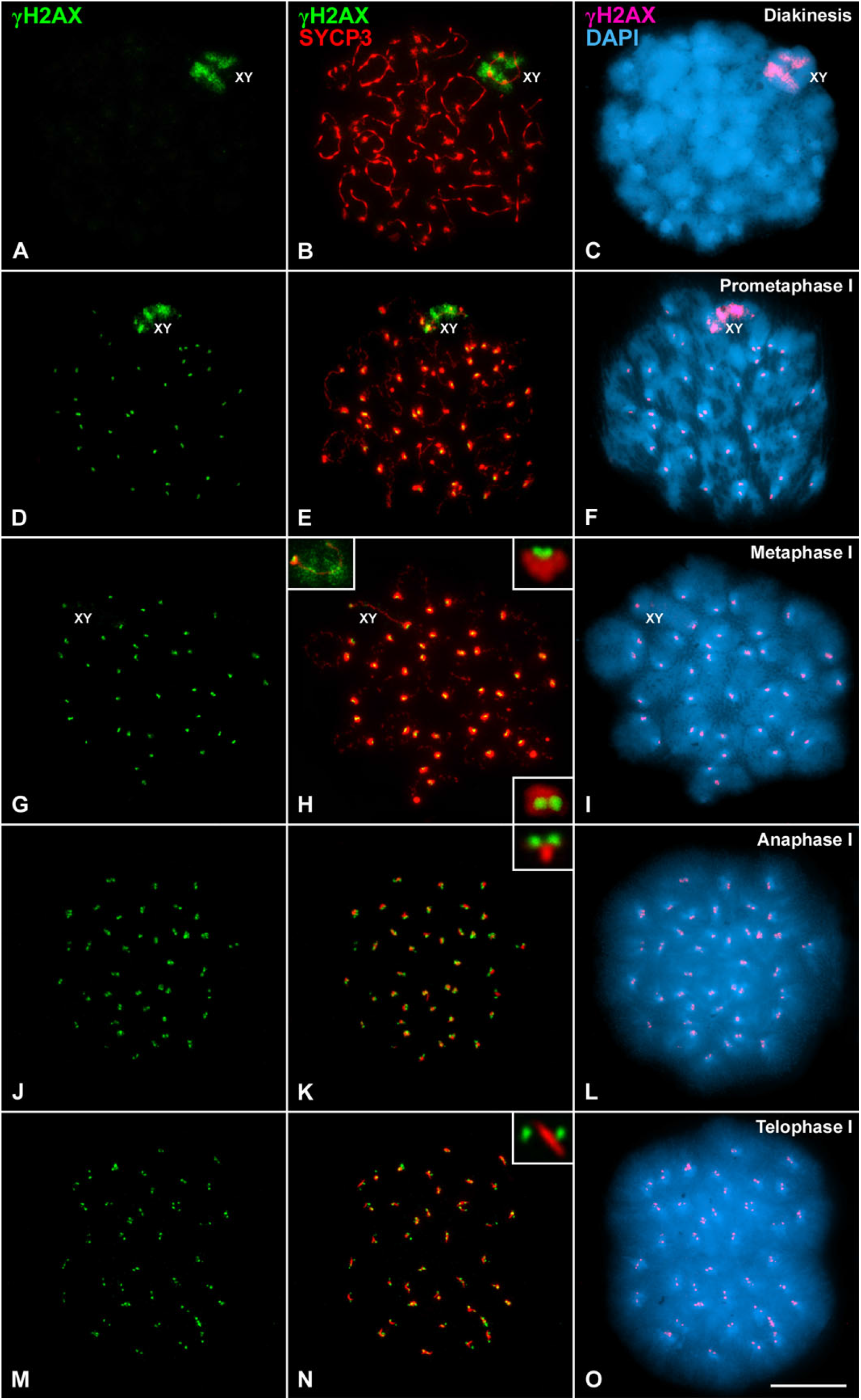
γ-H2AX distribution in post-diplotene stages of the first meiotic division. **(A, B, D, E, G, H, J, K, M, N)** Double immunolabeling of γ-H2AX (green) and SYCP3 (red). **(C, F, I, L, O)** Corresponding chromatin staining with DAPI (blue) and immunolabeling of γ-H2AX (pseudocoloured in pink). **(A-C)** Diakinesis, **(D-F)** prometaphase I, **(G-I)** metaphase I, **(J-L)** anaphase I and **(M-O)** telophase I spread spermatocytes. Left inset in **(H)** corresponds to a sex bivalent whose chromatin is γ-H2AX-labeled. Right insets in **(H)** show selected side and top views of a metaphase I centromere. Insets in **(K)** and **(N)** show representative views of anaphase I and telophase I centromeres. The position of sex bivalents (XY) is indicated. Details of centromeres in **(H, K, N)** correspond to a 400% magnification. Scale bar: 10 μm.

During anaphase I (Fig. 1J-L) and telophase I (Fig. 1M-O), SYCP3 remnants persisted at centromeres and the pairs of dots of γ-H2AX appeared more separated than during metaphase I (Fig. 1G-I). A detailed observation of centromeres showed SYCP3 bars between the pairs of γ-H2AX dots during anaphase I (Fig. 1K; inset) and telophase I (Fig. 1N; inset). In interkinesis spermatocytes SYCP3 appeared as elongated bars at chromocenters but no γ-H2AX signals were observed within the nuclei (Fig. 2A-C). During the second meiotic division γ-H2AX became detectable again (Fig. 2D-L). About twenty pairs of dots of γ-H2AX were visualized in prophase II nuclei (Fig. 2D-F) and in metaphase II spermatocytes (Fig. 2G-I). Those pairs of γ-H2AX signals located at the centromeric regions which are more brightly stained with DAPI (Fig. 2F and I). During anaphase II γ-H2AX signals became diffuse (Fig. 2J-L) and were not visualized in telophase II spermatocytes (data not shown) or early round spermatids (Fig. S2A and B). A burst of γ-H2AX labeling was again observed in mid elongated spermatids firstly visualized throughout the nuclei and that then decreased from the proximal area where the sperm flagella is assembled (Fig. S2C-F). However, γ-H2AX disappeared from late elongated spermatids (Fig. S2G and H). We also analyzed the distribution of γ-H2AX in spermatogonial metaphase chromosomes. In these chromosomes, γ-H2AX was detected as a diffuse labeling at some centromeres and at chromosome arms, but not as pairs of dots (Fig. S2I and J).

**Fig. 2.**
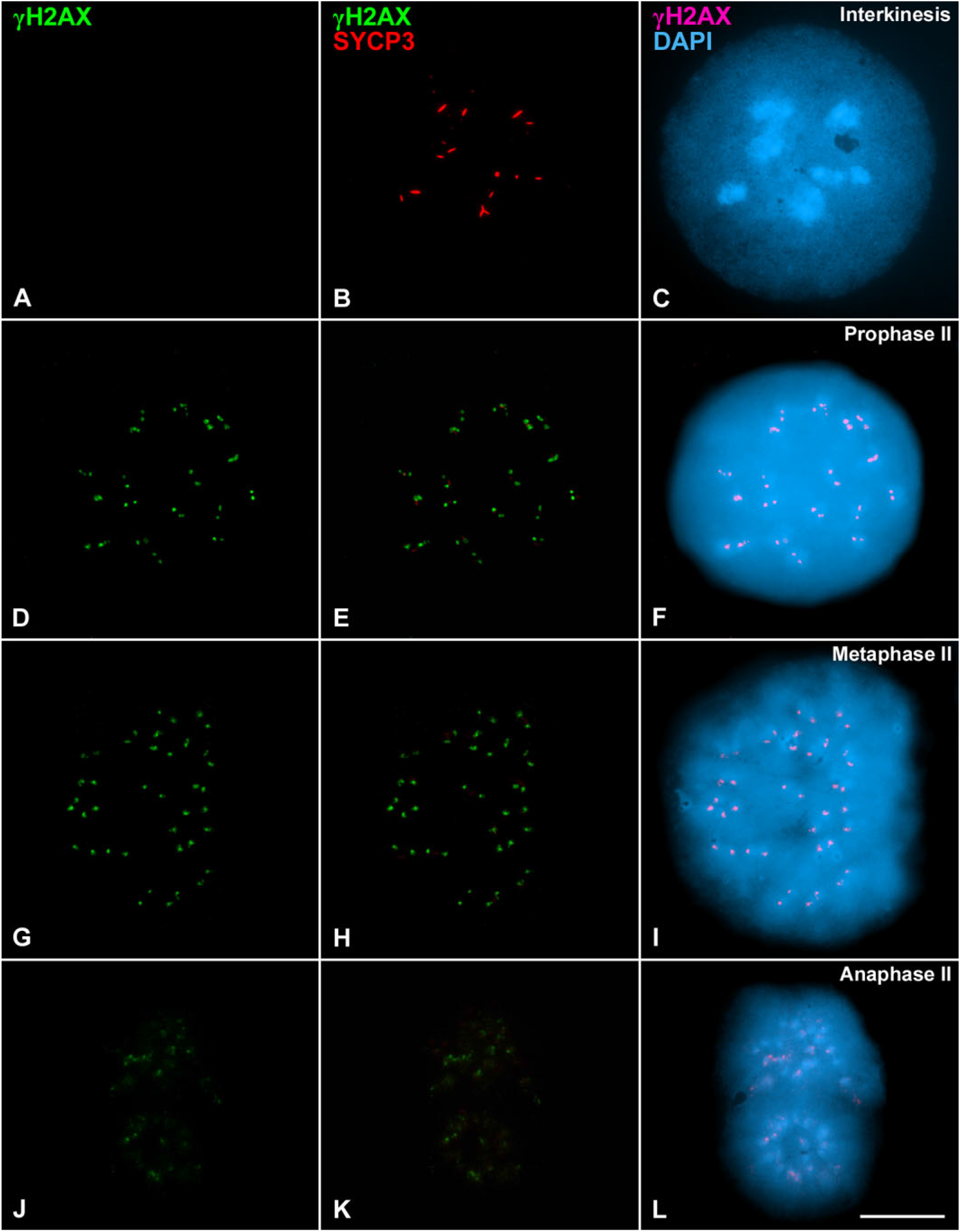
γ-H2AX distribution during the second meiotic division. **(A, B, D, E, G, H, J, K)** Double immunolabeling of γ-H2AX (green) and SYCP3 (red). **(C, F, I, L)** Corresponding chromatin staining with DAPI (blue) and immunolabeling of γ-H2AX (pseudocoloured in pink). **(A-C)** Interkinesis, **(D-F)** prophase II, **(G-I)** metaphase II and **(J-L)** anaphase II spread spermatocytes. Scale bar: 10 μm.

### γ-H2AX is present at the inner kinetochore plates

Altogether, our results suggested that γ-H2AX was located at kinetochores. To test this hypothesis we first made a double immunolabeling of γ-H2AX and H3K9me3, a post-translational modification of histone H3 that is a marker of centromeric heterochromatin. In metaphase I spermatocytes, H3K9me3 labeled intensely the centromeric heterochromatin leaving two unlabeled holes in which the two round signals of γ-H2AX were located (Fig. 3A and B; insets in both). A similar situation was found at anaphase I centromeres (Fig. 3C and D). Moreover, H3K9me3 marked the chromatin of the sex bivalent as γ-H2AX at metaphase I (Fig. 3A and B) and the X and Y chromatin at anaphase I (Fig. 3C and D).

**Fig. 3.**
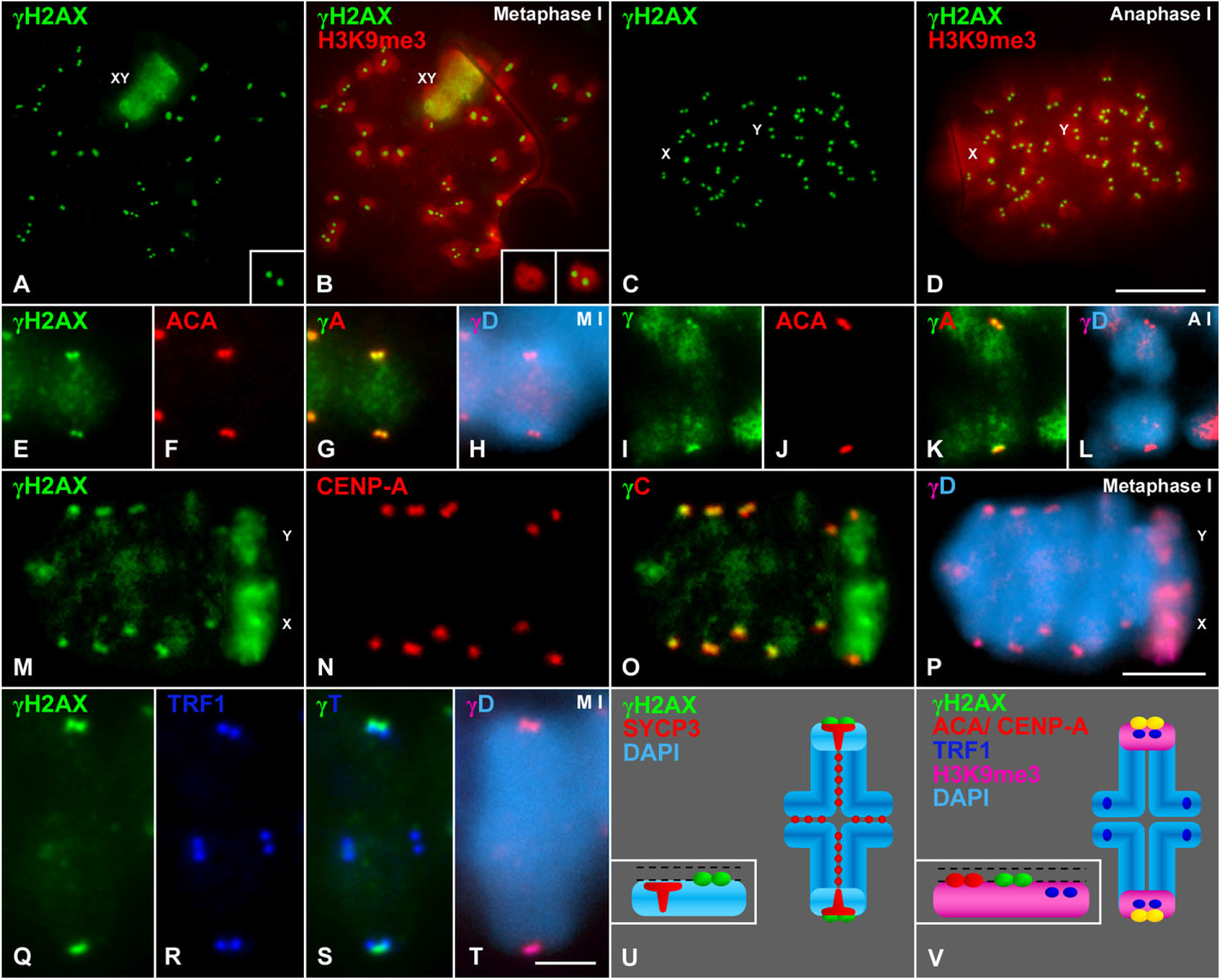
γ-H2AX signals at metaphase I and anaphase I centromeres in relation to H3K9me3, ACA, CENP-A and TRF1. **(A-D)** Double immunolabeling of γ-H2AX (green) and H3K9me3 (red) in **(A, B)** metaphase I and **(C, D)** anaphase I spread spermatocytes. Insets in **(A, B)** show an enlarged metaphase I centromere. **(E-L)** Double immunolabeling of γ-H2AX (green) and kinetochores by an ACA serum (red) in a selected autosomal metaphase I bivalent **(E-H)** and in segregating chromosomes in anaphase I **(I-L)** from squashed spermatocytes. **(M-P)** Double immunolabeling of γ-H2AX (green) and CENP-A (red) in a metaphase I squashed spermatocyte. Images show a partial projection of all focal planes. **(Q-T)** Double immunolabeling of γ-H2AX (green) and TRF1 (dark blue) in a selected autosomal metaphase I bivalent from a squashed spermatocyte. **(H, L, P, T)** Chromatin stained with DAPI (light blue) and corresponding immunolabeling of γ-H2AX (pseudocoloured in pink). **(U, V)** Schematic representations of the location of γ-H2AX, SYCP3, ACA, CENP-A, TRF1 and H3K9me3 signals in an autosomal metaphase I bivalent. Insets show the relative position of signals. Discontinuous black lines represent the outer and inner kinetochore plates. The position of sex bivalents (XY) and sex chromosomes (X, Y) are indicated. Details of the centromere included in **(A, B)** correspond to a 200% magnification. Scale bars: (A-D) 10 μm; (E-P) 10 μm; (Q-T) 5 μm.

To validate the results obtained on spread spermatocytes we employed the squashing technique that preserves the nuclear volume and chromosome condensation and positioning inside dividing spermatocytes (Page et al., 1998; Parra et al., 2002) in contrast to the drying-down spreading technique (Peters et al., 1997). Thus, the squashing technique allowed us to study the spatial relationships between γ-H2AX and other known kinetochore proteins. We first performed a double immunolabeling of γ-H2AX with an anti-centromere autoantibody (ACA) serum which labels kinetochores. The labeling of γ-H2AX obtained on squashed spermatocytes was the same as that observed on spread spermatocytes. In this sense, two pairs of intimately associated dots of γ-H2AX colocalizing with the sister kinetochore ACA signals were visualized at each autosomal metaphase I centromere (Fig. 3E-H). Likewise, γ-H2AX and kinetochore signals colocalized at the centromeres of segregating autosomal anaphase I chromosomes (Fig. 3I-L). Since the trilaminar mammalian kinetochores present outer and inner plates (Masatoshi and Fukagawa, 2020), we next investigated the precise localization of γ-H2AX signals at kinetochores. Since ACA sera recognize several kinetochore proteins, and to achieve a more precise location of γ-H2AX signals at kinetochores, we performed a double immunolabeling of γ-H2AX and CENP-A, a histone H3 variant specifically present at the inner kinetochore plates (Fig. 3M-P). CENP-A was visualized as two closely associated round signals representing the inner plates of both sister kinetochores at metaphase I centromeres (Fig. 3N), as previously described (Parra et al., 2004). These CENP-A signals colocalized with the γ-H2AX ones (Fig. 3O) suggesting that γ-H2AX is also present at, or very close to, the inner kinetochore plates. Since it has been previously described that proximal telomeres are just below sister kinetochores at metaphase I mouse centromeres (Viera et al., 2003), we then asked for the relative distribution of γ-H2AX and the telomeric protein TRF1 at metaphase I centromeres (Fig. 3Q-T). Our results showed that four pairs of TRF1 signals were detected at metaphase I bivalents, two pairs at distal telomere regions, and two pairs at proximal telomere regions close to centromeres (Fig. 3R and S). The superimposition of TRF1 and γ-H2AX signals at centromeres showed that proximal TRF1 signals were located just beneath the γ-H2AX ones (Fig. 3S). Altogether, our results support that γ-H2AX is present at the inner plates of both sister kinetochores (Fig. 3 U and V).

Finally, and since it has been described that the DDR kinase ATR is present at mitotic kinetochores (Kabeche et al., 2018), we also analyzed its possible presence at mouse meiotic kinetochores. Our results showed that ATR was present as numerous foci along synapsed and unsynapsed AEs in zygotene spermatocytes (Fig. S3A-C), representing early recombination nodules (Moens et al., 1999), but was absent from metaphase I or metaphase II centromeres or kinetochores (Fig. S3D-F).

## Discussion

In this study we show for the first time that γ-H2AX is present at the inner kinetochore plates during both male mouse meiotic divisions. In this regard, our results support the previous localization of γ-H2AX at the centromeres of metaphase I and anaphase I chromosomes from a grasshopper species (Cabrero et al., 2007). Accordingly, γ-H2AX appears as a structural phosphorylated histone variant that colocalizes cytologically with CENP-A, the canonical histone H3 variant present at the inner kinetochore plate, that confers epigenetic identity to the centromere and determines an accurate kinetochore assembly (McKinley and Cheeseman, 2016; Das et al., 2017).

### A novel role for γ-H2AX at kinetochores?

The chromatin present at kinetochores, called centrochromatin, presents interpersed nucleosomes with either CENP-A or histone H3 (McKinley and Cheeseman, 2016; Musacchio and Desai, 2017). It has been described that in CENP-A-containing nucleosomes this variant may be post-translationally modified by phosphorylation, acetylation, ubiquitination and methylation (Fukagawa, 2017; Srivastava and Foltz, 2018), while in histone H3-containing ones this histone may be methylated and acetylated at some residues (Srivastava et al., 2018). However, there are few studies reporting the presence and role of histone H2A post-translational modifications or variants at mammalian kinetochores. It has been proposed that at mammalian kinetochores histone H2A would be present at CENP-A containing nucleosomes, whereas the variant H2AZ would be present at histone H3-containing ones (Greaves et al., 2007). Regarding the histone variant H2AX, it has been demonstrated that in human cells the kinase Aurora-B phosphorylates its serine 121 (H2AXS121ph), that localized at the kinetochores, to promote the autophosphorylation of Aurora-B at mitotic centromeres (Shimada et al., 2016).

It has also been reported that during mitosis, and in the absence of DNA damage, γ-H2AX appears at kinetochores to recruit the DDR protein MDC1 that regulates the loading of the SAC proteins MAD2 and CDC20 to the kinetochores (Eliezer et al., 2014). Interestingly, the kinase ATR, a DDR protein, has also been reported at mitotic kinetochores (Kabeche et al., 2018). Our results show that γ-H2AX is present at mouse meiotic kinetochores, but not the kinase ATR. Thus, a possible relationship of γ-H2AX and the DDR pathway remains uncertain. However, γ-H2AX could have a specific role at mammalian kinetochores during meiosis. In this regard, γ-H2AX could collaborate in the recruitment of other proteins at kinetochores, as reported for γ-H2AX in mitotic kinetochores (Eliezer et al., 2014). Altogether, our results point that further studies are necessary to determine the possible participation of γ-H2AX in the kinetochore assembly and its putative role in chromosome segregation during mammalian meiosis.

## Materials and methods

### Animals and ethics statement

Wild-type C57BL/6 adult male mice were used in this study. Animals were handled according to the regulatory standards approved by the UAM Ethics Committee.

### Spreading of spermatocytes and squashing of seminiferous tubules

Testes from adult males were removed, detunicated, and seminiferous tubules then processed for spermatocyte spreading by the drying-down technique previously described (Peters et al., 1997), or processed for squashing as previously reported (Page et al., 1998; Parra et al., 2002).

### Immunofluorescence microscopy

Spread spermatocyte and squashed seminiferous tubules preparations were rinsed three times for 5 min in PBS and incubated in a humid chamber overnight at 4°C with the corresponding primary antibodies diluted in PBS. In double-immunolabeling experiments, primary antibodies from different host species were incubated simultaneously. Following three washes in PBS for 5 min, the slides were incubated for 1 h at room temperature with the corresponding secondary antibodies. The slides were subsequently rinsed in PBS and counterstained for 3 min with 10 μg/ml DAPI (4’,6-diamidino-2-phenylindole). After a final rinse in PBS, the slides were mounted with Vectashield (Vector Laboratories) and sealed with nail polish.

### Antibodies for immunofluorescence

We used three different antibodies against γ-H2AX: a mouse monoclonal antibody (Millipore, 05-636) at a 1:500 dilution, and two rabbit polyclonal antibodies (Abcam, ab-2893; Cell Signaling, 2577S) used at 1:100 and 1:10 dilutions, respectively. The three antibodies used in this study offered the same staining patterns in immunofluorescence. Other primary antibodies were used as follows: a mouse monoclonal antibody against mouse SYCP3 (Santa Cruz, sc-74569) at 1:100; a rabbit polyclonal antibody against human SYCP3 (Abcam, ab-150292) at 1:100; a rabbit polyclonal antibody against H3K9me3 (Abcam, ab-8898) at 1:100; a human anti-centromere autoantibody (ACA serum) labeling kinetochores (Antibodies Incorporated, 435-2RG-7) at 1:20; a rabbit polyclonal antibody against mouse TRF1 (Alpha Diagnostic, TRF12-S) at 1:100; the monospecific rabbit polyclonal serum SS2 raised against a synthetic peptide covering amino acid residues 3-17 of human CENP-A (Valdivia et al., 1998) at 1:50; and a goat polyclonal antibody against human ATR (Santa Cruz, sc-1887) at 1:10. The following secondary antibodies were employed at a 1:100 dilution: Alexa 488-conjugated donkey anti-mouse IgG (Molecular Probes, A-21202), Alexa 594-conjugated donkey anti-mouse IgG (Molecular Probes, A-21203), Alexa 488-conjugated donkey anti-rabbit IgG (Molecular Probes, A-21206), Texas Red-conjugated donkey anti-rabbit IgG (Jackson, 711-075-152), Texas Red-conjugated donkey anti-human IgG (Jackson, 709-075-149), and Alexa 488-conjugated donkey anti-goat IgG (Molecular Probes, A-11055).

### Image acquisition

Observations were performed using an Olympus BX61 microscope equipped with a motorized Z axis and epifluorescence optics. Images were captured with an Olympus DP71 digital camera controlled by the CellSens Dimension software and processed with Adobe Photoshop.

## Abbreviations used in this paper

ACA: anti-centromere autoantibody
DDR: DNA damage response
DSBs: DNA double-strand breaks
SAC: spindle-assembly checkpoint

## Acknowledgments

This work was supported by grant BFU2014-53681-P from Ministerio de Economía y Competitividad (Spain). AG was supported by “Ayuda para el Fomento de la Investigación en Estudios de Máster”, and “Ayudas PostMáster del Departamento de Biología” from Universidad Autónoma de Madrid.

## Author contributions

AV and JAS conceived the study; MMV obtained the anti-CENP-A antibody; AG, AV, MTP, JSR and JAS performed all experiments, analyzed results and wrote the paper.

## Supplementary figure legends

**Fig. S1.**
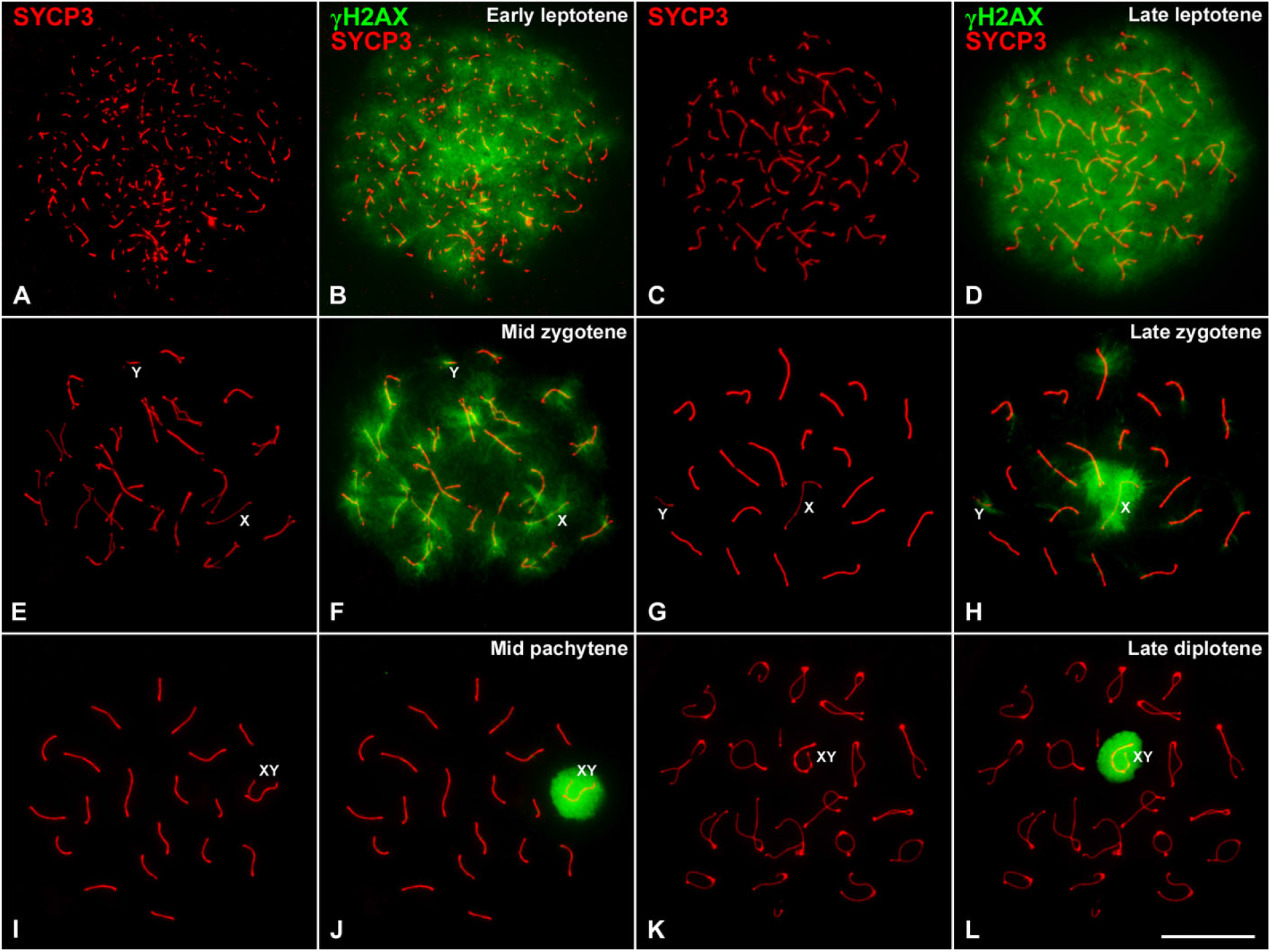
γ-H2AX distribution during prophase I stages. **(A-L)** Double immunolabeling of γ-H2AX (green) and SYCP3 (red) in **(A, B)** early leptotene, **(C, D)** late leptotene, **(E, F)** mid zygotene, **(G, H)** late zygotene, **(I, J)** mid pachytene and **(K, L)** late diplotene spread spermatocytes. The position of sex chromosomes (X, Y) and bivalents (XY) is indicated. Scale bar: 10 μm

**Fig. S2.**
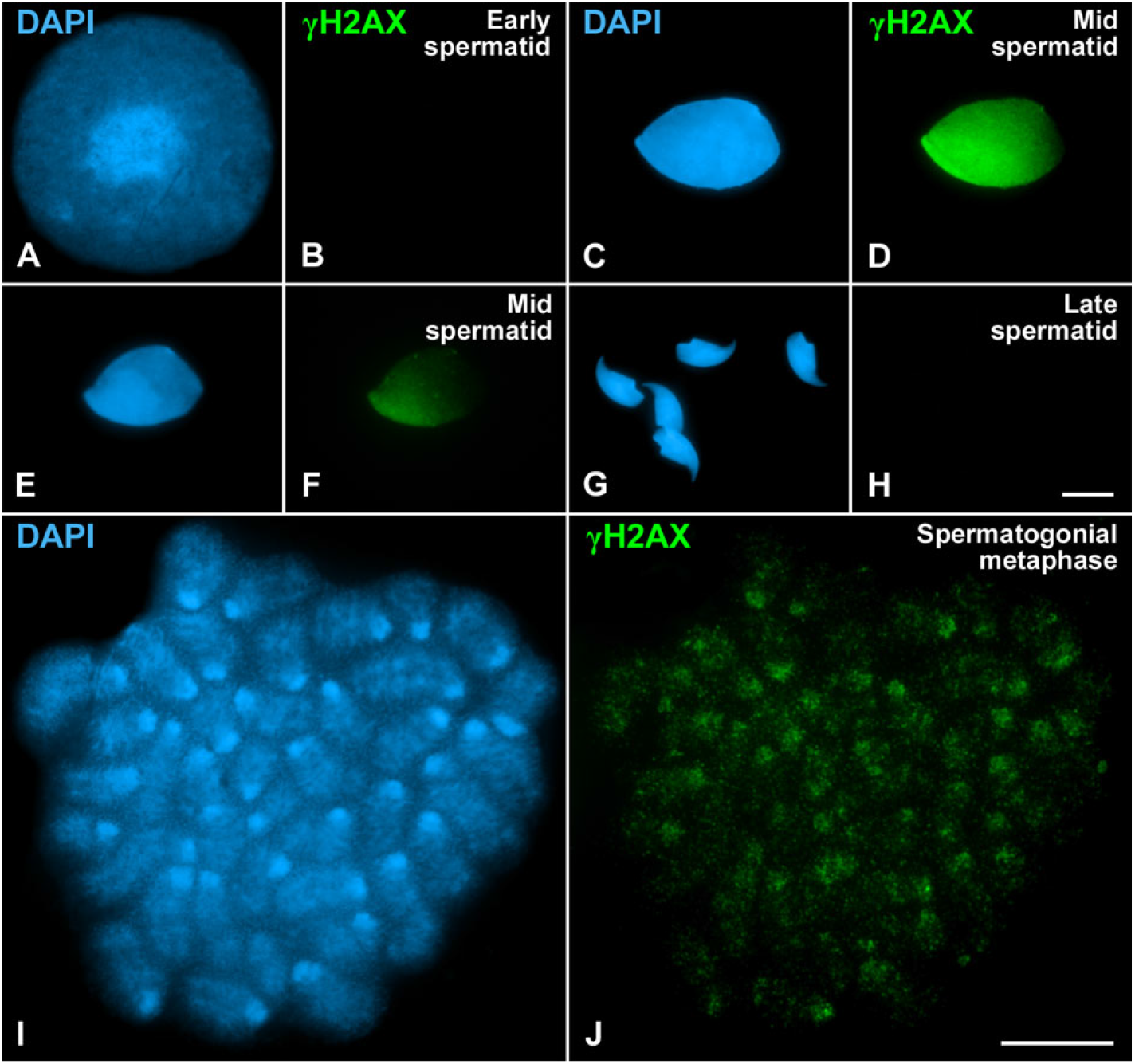
γ-H2AX distribution during spermiogenesis and spermatogonial metaphase. Immunolabeling of γ-H2AX (green) and DAPI staining of chromatin (blue) in **(A, B)** early round spermatid, **(C-F)** mid elongated spermatids, **(G, H)** late spermatids and **(I, J)** spermatogonial metaphase. Scale bars: (A-H) 10 μm; (I and J) 10 μm.

**Fig. S3.**
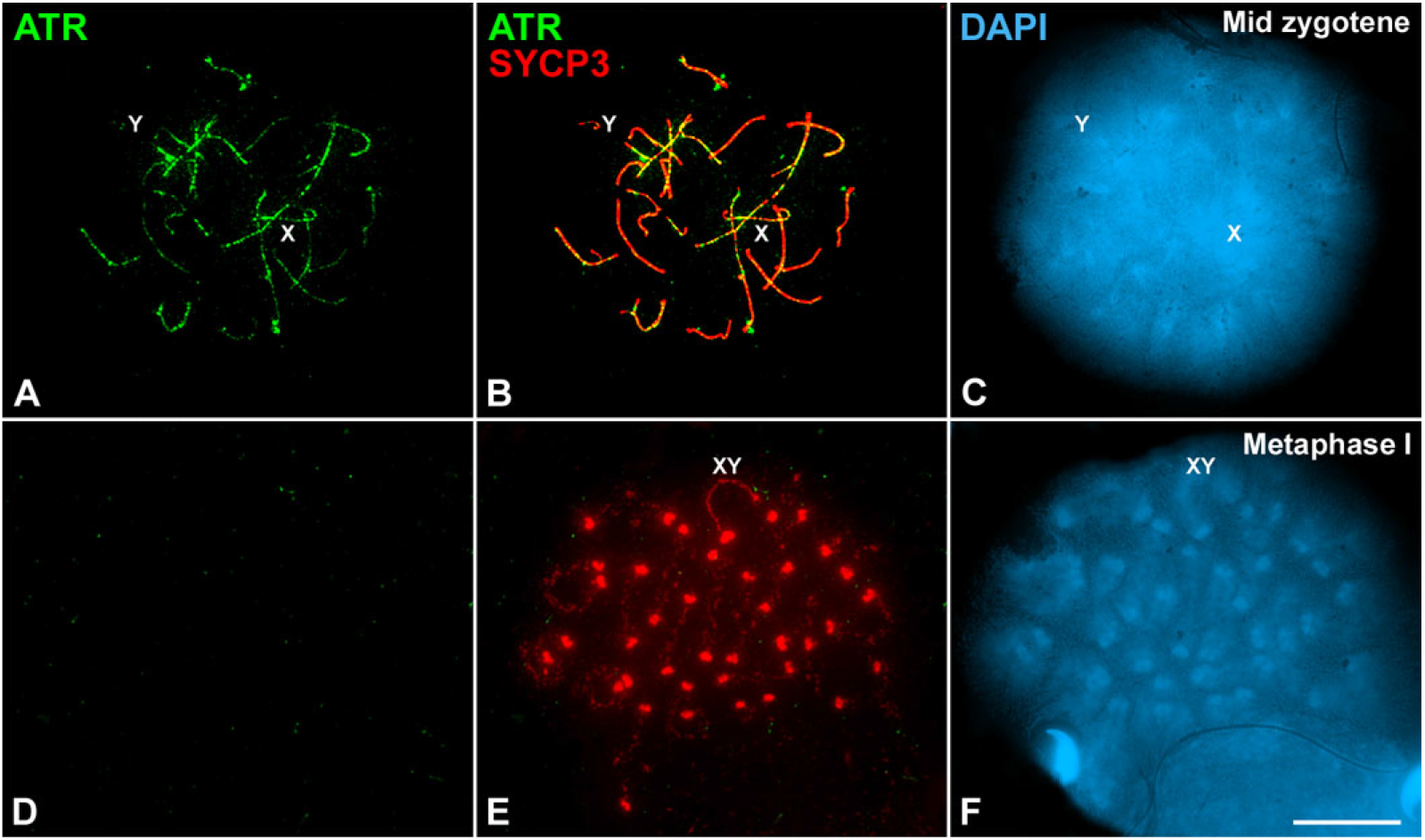
ATR distribution in zygotene and metaphase I spermatocytes. **(A-F)** Double immunolabeling of ATR (green) and SYCP3 (red) and DAPI staining of chromatin (blue) in **(A-C)** zygotene and **(D-F)** metaphase I spread spermatocytes. The position of sex chromosomes (X, Y) and bivalent (XY) is indicated. Scale bar: 10 μm.

